# SLC45A2 protein stability and regulation of melanosome pH determine melanocyte pigmentation

**DOI:** 10.1101/2020.03.03.974881

**Authors:** Linh Le, Iliana E. Escobar, Tina Ho, Ariel J. Lefkovith, Emily Latteri, Megan K. Dennis, Kirk D. Haltaufderhyde, Elena V. Sviderskaya, Dorothy C. Bennett, Elena Oancea, Michael S. Marks

## Abstract

*SLC45A2* encodes a putative transporter expressed primarily in pigment cells. *SLC45A2* mutations and polymorphisms cause oculocutaneous albinism (OCA) and pigmentation variation, but neither SLC45A2 localization and function nor how gene variants affect these properties are known. We show that SLC45A2 localizes to mature melanosomes that only partially overlap with a cohort expressing the chloride channel OCA2. SLC45A2 expressed ectopically in HeLa cells localizes to lysosomes and raises lysosomal pH, suggesting that, like OCA2, SLC45A2 in melanocytes de-acidifies maturing melanosomes to support melanin synthesis. Analyses of SLC45A2- and OCA2-deficient mouse melanocytes show that SLC45A2 functions later during melanosome maturation than OCA2, and that OCA2 overexpression compensates for loss of SLC45A2 expression in pigmentation. The light skin-associated SLC45A2 allelic F374 variant restores only moderate pigmentation to SLC45A2-deficient melanocytes because of low level expression in melanosomes due to rapid proteasome-independent degradation. Our data indicate that SLC45A2 maintains melanosome neutralization – initially orchestrated by transient OCA2 activity – to support melanization at late stages of melanosome maturation, and that a common variant imparts reduced activity due to protein instability.

## INTRODUCTION

Melanins are the main source of pigmentation in the skin, hair, and eyes of mammals and other vertebrates. In humans, melanins serve as a barrier to the harmful effects of ultraviolet radiation and play an important role in the development and functioning of the retina (d’Ischia, Wakamatsu et al., 2015). Melanin synthesis takes place in skin and ocular melanocytes and in ocular pigment epithelia within specialized organelles called melanosomes (Hearing, 2005, Marks & Seabra, 2001, Seiji, Fitzpatrick et al., 1963).

Heritable defects in melanin synthesis underlie the various forms of oculocutaneous albinism (OCA), characterized by impaired vision and increased susceptibility to skin and ocular cancers (Montoliu, Grønskov et al., 2014). To date, non-syndromic OCA has been linked to inactivating mutations in seven different genes (Montoliu et al., 2014).

Sequence variation at the loci for some of these genes has been linked to variability in skin, hair and eye color among humans (Adhikari, Mendoza-Revilla et al., 2019, Branicki, Brudnik et al., 2008, Crawford, Kelly et al., 2017, Han, Kraft et al., 2008, Lamason, Mohideen et al., 2005, Liu, Visser et al., 2015, Martin, Lin et al., 2017, Stokowski, Pant et al., 2007). Nevertheless, the molecular function of the majority of the OCA genes has not yet been fully characterized.

OCA type 4 (OMIM #606574) represents 3-12% of total OCA patients in population studies (Gronskov, Ek et al., 2009, Lasseaux, Plaisant et al., 2018, Mauri, Manfredini et al., 2017, Wei, Wang et al., 2010, Wei, Zang et al., 2015) and is due to mutations in the *SLC45A2* gene encoding the putative transmembrane transporter SLC45A2 (a.k.a. membrane associated transporter protein, MATP or antigen isolated from immuno-selected melanoma-1, AIM1)(Newton, Cohen-Barak et al., 2001). Mutations in the homologous gene underlie pigment dilution in a number of vertebrate species, including gorilla, several breeds of dog, tigers, horses, mice, shrew, chickens, pigeons, quail, frogs, fish and perhaps cattle (Caduff, Bauer et al., 2017, DeLay, Corkins et al., 2018, Domyan, Guernsey et al., 2014, Dooley, Schwarz et al., 2013, Fukamachi, Shimada et al., 2001, Gunnarsson, Hellström et al., 2007, Mariat, Taourit et al., 2003, Minvielle, Cecchi et al., 2009, Newton et al., 2001, Prado-Martinez, Hernando-Herraez et al., 2013, Rothammer, Kunz et al., 2017, Tsetskhladze, Canfield et al., 2012, Tsuboi, Hayashi et al., 2009, Wijesena & Schmutz, 2015, Winkler, Gornik et al., 2014, Xu, Dong et al., 2013), and polymorphisms at the *SLC45A2* locus are associated with skin tone differences and skin aging in several human population studies (Adhikari et al., 2019, Branicki et al., 2008, Cerqueira, Hunemeier et al., 2014, Fracasso, de Andrade et al., 2017, Han et al., 2008, Jonnalagadda, Norton et al., 2016, Law, Medland et al., 2017, Liu et al., 2015, Lopez, Garcia et al., 2014, Soejima & Koda, 2007, Stokowski et al., 2007, Yuasa, Umetsu et al., 2006). OCA4 patients have very low levels of pigmentation and phenotypically resemble OCA2 patients who lack the melanosomal chloride channel, OCA2 (Bellono, Escobar et al., 2014), suggesting that SLC45A2 plays an important role in melanogenesis (Montoliu et al., 2014). Moreover, primary melanocytes from mice carrying the inactivating *underwhite* (uw) mutation of *Slc45a2* (Du & Fisher, 2002, Newton et al., 2001) are severely hypopigmented (Costin, Valencia et al., 2003), indicating a melanocyte-intrinsic defect; this would be consistent with the known restriction of SLC45A2 expression to pigment cells and a few other cell types (Baxter & Pavan, 2002, Bin, Bhin et al., 2015, Harada, Li et al., 2001, Loftus, Larson et al., 2002). However, while the 12-transmembrane domain SLC45A2 protein bears weak homology to sucrose transporters in plants and *Drosophila* (Lemoine, 2000, Meyer, Vitavska et al., 2011, Newton et al., 2001), its function in melanocytes is not understood.

When expressed in yeast, mouse SLC45A2 functions at the plasma membrane as an acid-dependent importer of sugars (sucrose, glucose, or fructose) into the cytosol (Bartölke, Heinisch et al., 2014), suggesting that if SLC45A2 localized to acidic organelles it might facilitate export of a sugar and protons from the lumen to the cytosol. Neutralization of acidic early stage melanosomes is a critical process for melanogenesis (Bellono et al., 2014, Raposo, Tenza et al., 2001), as the key enzyme in melanogenesis, Tyrosinase, is inactive at pH < 6 (Ancans, Tobin et al., 2001, Halaban, Patton et al., 2002). Consistent with a function in proton export and neutralization of acidic organelles, pigmentation of a zebrafish *Slc45a2* mutant was rescued upon inhibition of endolysosomal and melanosomal acidification by treatment with bafilomycin A1 or by knockdown of the atp6v1 subunit of the vacuolar ATPase (Dooley et al., 2013), and knockdown of SLC45A2 in a pigmented melanoma cell line resulted in increased acidification of early stage melanosomes (Bin et al., 2015). However, despite limited evidence for SLC45A2 on melanosomes (Bin et al., 2015), the localization of SLC45A2 within melanocytes has not yet been firmly established and it is not clear whether the effects of *SLC45A2* mutants reflect a direct effect of SLC45A2 function on the lumenal environment of melanosomes or an indirect effect due to impaired function of other organelles during melanosome maturation, as appears to be the case for the endolysosomal transporter MFSD12 (Crawford et al., 2017). Moreover, if SLC45A2 indeed functions to neutralize melanosome pH, its function must be coordinated with that of OCA2, a major regulator of melanosomal pH (Bellono et al., 2014). While mice lacking expression of either SLC45A2 or OCA2 each have dramatic coat color dilution (Dickie, 1964, Sweet, Brilliant et al., 1998), mice with hypomorphic mutations in both *Slc45a2* and *Oca2* are more severely hypopigmented than either mutant alone (Lehman, Silvers et al., 2000), suggesting that the encoded proteins have distinct functions.

Differences in skin and hair pigmentation among European, Chinese, South American and South Asian human populations have been ascribed to a single genetic variant in SLC45A2 associated with the SNP rs16891982. This encodes a single amino acid change at position 374 – leucine (L374) in dark skinned individuals and phenylalanine (F374) in light skinned individuals (Adhikari et al., 2019, Branicki et al., 2008, Cerqueira et al., 2014, Han et al., 2008, Jonnalagadda et al., 2016, Liu et al., 2015, Lopez et al., 2014, Soejima & Koda, 2007, Stokowski et al., 2007, Yuasa et al., 2006); this residue lies within the 8th predicted transmembrane domain of SLC45A2 (Newton et al., 2001). The light F374 variant is nearly fixed in light-skinned human populations (Soejima & Koda, 2007). When introduced into a plant sucrose transporter, the amino acid corresponding to F374 resulted in a 90% decrease in transporter activity without influencing the affinity for substrate (Reinders & Ward, 2015), and slc45a2 with an introduced F374 mutation was unable to rescue pigmentation in a *slc45a2* mutant zebrafish (Tsetskhladze et al., 2012). However, what mechanisms underlie the decreased activity of the F374 variant is not understood.

Analyses of SLC45A2 localization and function in pigment cells have been hindered by the lack of suitable specific antibodies. Here we assess the localization and function of SLC45A2 L374 and F374 variants in melanosome biogenesis by analyzing epitope-tagged human SLC45A2 expressed in wild-type melanocytes, *Slc45a2*-deficient melanocytes from *underwhite* mice, and HeLa cells, and by comparing the phenotypes of *underwhite* and OCA2-deficient *pink-eyed dilute* melanocytes. We show that OCA2 and SLC45A2 directly neutralize the melanosome matrix at distinct steps during maturation, and that the F374 variation accelerates SLC45A2 degradation but does not alter its localization. Our data indicate that melanosome neutralization is a critical process for the maintenance of eumelanin pigmentation that is regulated at more than one step during melanosome maturation.

## RESULTS

### Functionality of HA-SLC45A2

Available antibodies to SLC45A2 were not suitable in our hands for immunolocalization analyses. Therefore, we used epitope-tagging to investigate SLC45A2 localization in melanocytes. Human SLC45A2 was tagged at the N- or C-terminus with either the HA11 epitope tag or EGFP and cloned into a retroviral vector. To determine whether the tagged transgenes were functional, they were expressed by recombinant retroviral infection in *Slc45a2*-deficient melan-uw cells from *underwhite* mice (Dickie, 1964). The *Slc45a2^uw^* allele encodes a non-functional truncated SLC45A2 protein and an undetectable transcript (Du & Fisher, 2002, Newton et al., 2001). Consequently, *underwhite* mice (Dickie, 1964, Lehman et al., 2000, Sweet et al., 1998) and primary melanocytes derived from them (Costin et al., 2003) are severely hypopigmented. Likewise, melan-uw cells are very pale compared to “wild-type” (WT) immortalized melanocytes (melan-Ink4a) from C57BL/6-*Ink4a^-/-^* mice (Sviderskaya, Hill et al., 2002) (**Figure 1a, b**). While expression of the EGFP-tagged proteins failed to consistently restore pigmentation in melan-uw cells (data not shown), expression of the N-terminally HA-tagged human SLC45A2 (HA-SLC45A2) restored partial or full pigmentation in a substantial fraction of cells by 48-72 h post-infection, whereas pigmentation was not restored by expression of a similarly tagged unrelated polytopic protein - the putative sugar-nucleotide transporter, SLC35D3, which plays no role in pigmentation (Chintala, Tan et al., 2007) (**Figure 1c-g**). Additionally, quantitative analysis of melanin content in melan-uw cells stably expressing HA-SLC45A2 showed pigmentation at levels comparable to those of melan-Ink4a cells (see **Figure 5f**). These data indicate that HA-SLC45A2 is fully functional in melanocytes, and therefore validate its use in defining the localization and biosynthesis of SLC45A2.

**Figure 1.**
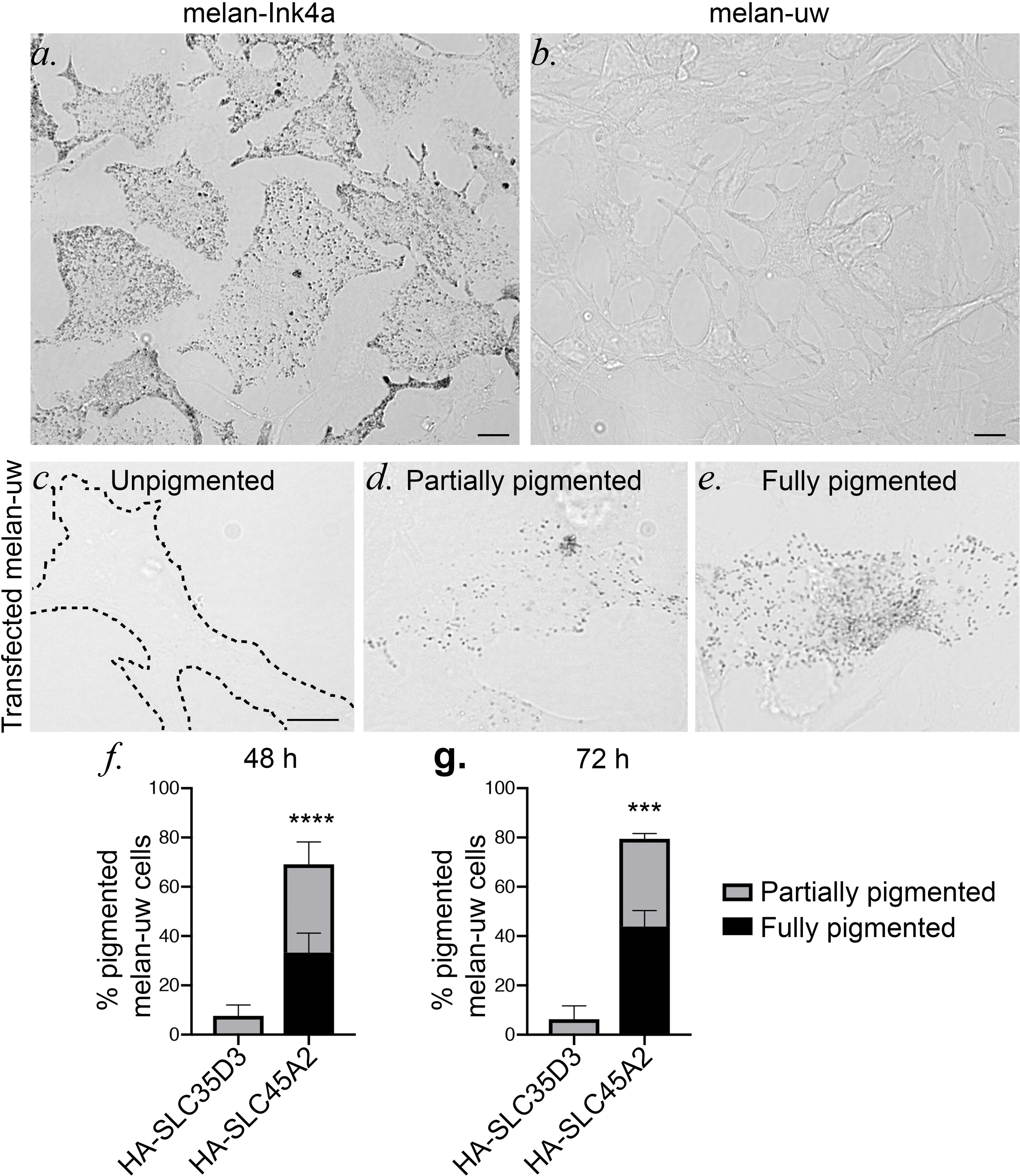
HA-SLC45A2 functionally restores pigmentation in *Slc45a2*-deficient *underwhite* melanocytes. Wide field microscopy with bright field illumination of melan-Ink4a (*a*) and melan-uw (*b*) cells to visualize pigmentation. Scale bar, 10 μm. Mouse melan-uw cells were transiently transfected with plasmids encoding human SLC45A2 with an N-terminal HA epitope (HA-SLC45A2) or with a comparably HA-tagged control polytopic protein, SLC35D3 (HA-SLC35D3). 48-72 h after transfection, HA-expressing cells were scored for their level of pigmentation. *c*-*e*, examples of unpigmented (*c*, like untransfected cells), partially pigmented (*d*) and fully pigmented (*e*) cells. Scale bar, 10μm. *f*, *g*, quantification of the percentage of HA-expressing cells with partial (gray) or full (black) pigmentation at 48 (*f*) or 72 (*g*) h after transfection. Data represent mean ± SD from 3 experiments; n = 234 (HA-SLC45A2) and 109 (HA-SLC35D3) at 48 h, 173 (HA-SLC45A2) and 108 (HA-SLC35D3) at 72 h. ***, p < 0.005; ****, p<0.0001 by unpaired two-tailed t-test.

### HA-SLC45A2 localizes to the limiting membrane of pigmented melanosomes and is partially enriched in a membrane subdomain

To define SLC45A2 localization in melanocytes, HA-SLC45A2 was expressed either stably from recombinant retroviruses in melan-uw cells or by transient transfection in WT melan-Ink4a melanocytes. Cells were then fixed and processed for immunofluorescence microscopy with image deconvolution (dIFM) using antibodies to the HA tag and to endogenous markers. In *Slc45a2*-deficient melan-uw cells, HA-SLC45A2 localized primarily to pigment granules (**Figure 2a, b, d**) often in a ring pattern around the granule (**Figure 2b**), in two to three puncta associated with the granule exterior (**Figure 2a**), or both (**Figure 2c**). The distinct patterns did not correlate with SLC45A2 expression levels. The melanosomes labeled by HA-SLC45A2 overlapped with those harboring the melanosomal enzymes Tyrosinase (TYR) and Tyrosinase-related protein-1 (TYRP1) (**Figure 2a, b, d**; overlap was 82.7 ± 1.3% with TYR and 79.2 ± 2.5% with TYRP1). By contrast, HA-SLC45A2-labeled structures overlapped poorly with LAMP2, a membrane protein of lysosomes but not melanosomes, which are distinct organelles in eumelanin-generating melanocytes (Raposo et al., 2001) (**Figure 2c, d**; overlap was 34.4% ± 2.9% with LAMP2). Essentially identical results were observed for HA-SLC45A2 transiently expressed in melan-Ink4a cells (**Figure EV1**), which express endogenous SLC45A2.

**Figure 2.**
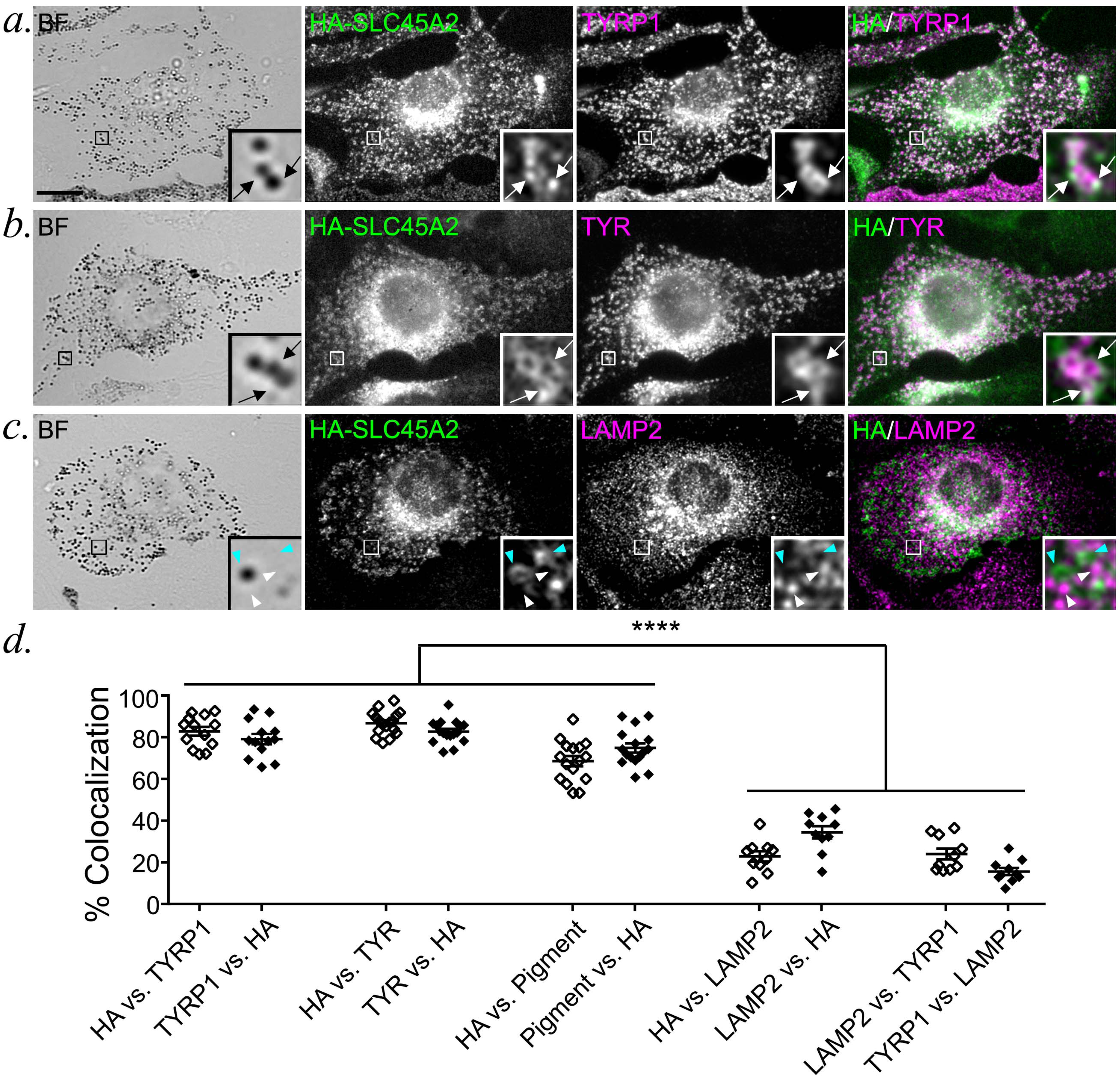
HA-SLC45A2 localizes to melanosomes when expressed in SLC45A2-deficient melan-uw melanocytes. (*a*-*c*) Melan-uw cells stably expressing HA-tagged SLC45A2 were fixed, labeled for HA (green) and for TYRP1 (*a*), TYR (*b*), or LAMP2 (*c*) (magenta), and analyzed by dIFM and bright field (BF) microscopy to visualize melanin. Insets of boxed regions are magnified 7.5 times. Scale bar, 10 μm. SLC45A2 that localized to TYRP1- or TYR-labeled compartments (white arrows) or SLC45A2 (cyan arrowheads) and LAMP2 (white arrowheads) on separate compartments are indicated. (d) The percentage of compartments with both markers in *a*-*c* was quantified by manual counting of at least 13 cells each from 3 separate experiments. Colocalization is represented as mean ± SEM of label 1 vs label 2, in which the number of compartments containing both label 1 and label 2 is presented as a percentage of the total number of compartments containing label 2. The % colocalization between TYRP1 and LAMP2 was quantified as a negative control. Statistical significance was determined using one-way ANOVA with Sidak’s test for multiple comparisons; only significant differences are indicated. ****, p < 0.001.

The degree of HA-SLC45A2 labeling on punctate subdomains vs. the melanosome limiting membrane was similar in transiently transduced melan-Ink4a and stably transduced melan-uw cells. Together, these data indicate that SLC45A2 localizes to mature melanosomes, where it is often enriched in subdomains associated with the limiting membrane.

### SLC45A2 localizes to lysosomes and regulates lysosomal pH when ectopically expressed in HeLa cells

When expressed in non-melanocytic cells, many melanosomal proteins localize to late endosomes and lysosomes (Ambrosio, Boyle et al., 2016, Bellono, Escobar et al., 2016, Berson, Harper et al., 2001, Bouchard, Fuller et al., 1989, Calvo, Frank et al., 1999, Piccirillo, Palmisano et al., 2006, Simmen, Schmidt et al., 1999, Sitaram, Piccirillo et al., 2009, Vijayasaradhi, Xu et al., 1995). Indeed, when expressed in HeLa cells and analyzed by dIFM, HA-SLC45A2 was detected on LAMP1-containing lysosomes – interestingly, in distinct punctate microdomains associated with the limiting membrane (**Fig. 3a**), similar to the distribution in some melanosomes in melanocytes. This indicates that SLC45A2 localizes to lysosomes when expressed ectopically in HeLa cells and retains its ability to accumulate in a membrane subdomain.

**Figure 3.**
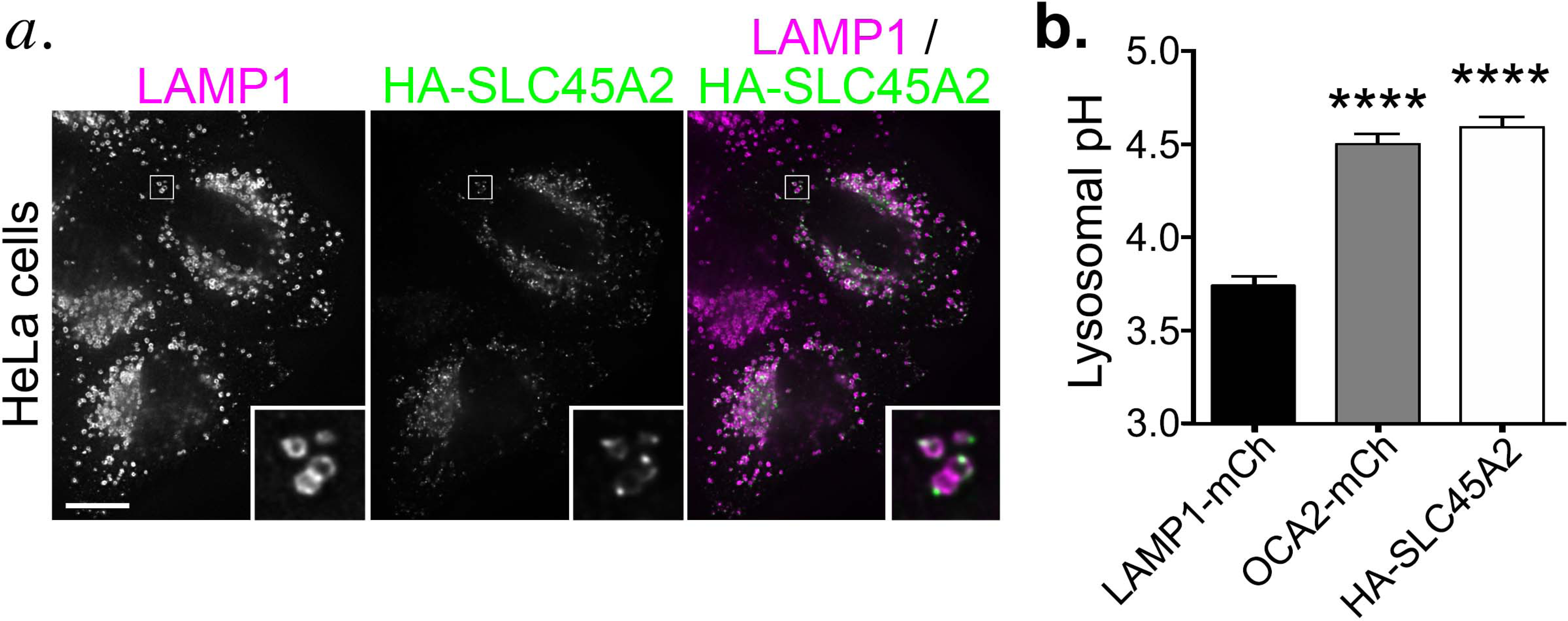
HA-SLC45A2 expressed in HeLa cells localizes to lysosomes and neutralizes lysosomal pH. *a*. HeLa cells transiently transfected with HA-SLC45A2 were fixed, labeled for HA (green) and the lysosomal protein LAMP1 (magenta), and analyzed by dIFM. Merged image is at right. Insets of boxes are magnified 5-fold. Scale bar, 10 μm. *b.* HeLa cells transiently transfected with LAMP1-mCherry alone, mCherry-OCA2, or HA-SLC45A2 and LAMP1-mCherry were incubated with Lysosensor DND-160, and analyzed by fluorescence microscopy with excitation wavelength 405. The W1/W2 emission ratio for mCherry-labeled endolysosomes was calculated and compared to that from a standard curve to define pH as described in Materials and Methods. Data represent mean values from at least 299 endolysosomes over 3 separate experiments. Statistical significance was determined using one-way ANOVA with Sidak’s test for multiple comparisons; only significant differences are indicated. ****, p < 0.001.

SLC45A2 was predicted to be a transmembrane transporter that influences melanosome pH (Bin et al., 2015, Newton et al., 2001). We have been unable to directly assess melanosomal pH in melanocytes using common cellular probes for lysosomal pH such as Lysotracker and Lysosensor because in our hands melanocytes fail to accumulate these probes intracellularly. However, we previously showed that the OCA2 chloride channel expressed in HeLa cells localizes to lysosomes and neutralizes their lumenal pH, paralleling its role in neutralizing the lumenal pH of melanosomes to enhance TYR activity and melanogenesis in melanocytes (Bellono et al., 2014). We therefore tested whether SLC45A2 expression affects lysosomal acidification in HeLa cells using the pH-sensitive ratiometric dye Lysosensor DND-160. HeLa cells were transfected with either LAMP1-mCherry alone as a negative control, mCherry-OCA2 as a positive control, or LAMP1-mCherry and HA-SLC45A2 together to identify HA-SLC45A2-expressing cells by live imaging (cotransfection efficiency measured by immunostaining was >90%). Cells preincubated with Lysosensor DND-160 were imaged at two emission wavelengths, and an average emission ratio was calculated for individual mCherry-LAMP1-positive compartments (see Materials and Methods). The corresponding pH for the average emission ratio was determined using a pH curve generated in each experiment by incubation in buffers with different pH values. Our results show that the average pH of lysosomes containing HA-SLC45A2 was nearly one pH unit higher than the pH of control cells, similar to the pH of mCherry-OCA2-containing lysosomes (**Fig. 3b**). These data suggest that, like OCA2, SLC45A2 can function to raise melanosomal pH.

### SLC45A2 and OCA2 function at distinct stages of melanosome maturation

The similar function of SLC45A2 and OCA2 in neutralizing organellar pH led us to ask whether these transporters function at the same or distinct stages of melanosome maturation. We first used dIFM to compare the distribution of HA-SLC45A2 and HA-OCA2 to melanosomes labeled by the mature melanosome marker TYRP1 in transiently transfected melan-Ink4a cells. HA-SLC45A2 and TYRP1 were largely present in the same compartments (86.58 ± 1.25% of HA-SLC45A2 in TYRP1-containing structures, and 89.17 ± 1.32% of TYRP1 in HA-SLC45A2-containing structures; **Figure 4a, e**), and the fluorescence signal intensity within positive compartments was highly correlated (**Figure 4c**). By contrast, although TYRP1 and OCA2 each primarily localize to subsets of pigment granules (Raposo et al., 2001, Sitaram, Dennis et al., 2012, Sitaram et al., 2009, Vijayasaradhi, Doskoch et al., 1991, Vijayasaradhi et al., 1995), they label only partially overlapping populations (58.57 ± 4.87% of HA-OCA2 in TYRP1-containing compartments, and 48.66 ± 5.32% of TYRP1 in OCA2-containing compartments; **Figure 4b, e**). Moreover, within those overlapping compartments, the fluorescence signal intensities of HA-OCA2 and TYRP1 over background correlated substantially less well than those of HA-SLC45A2 and TYRP1 (**Figure 4d**). These data suggest that OCA2 and SLC45A2 are enriched in melanosomes of distinct maturation stages, with SLC45A2 present in largely the same subset of mature melanosomes as TYRP1, while OCA2 occupies a partially different subset of melanosomes.

**Figure 4.**
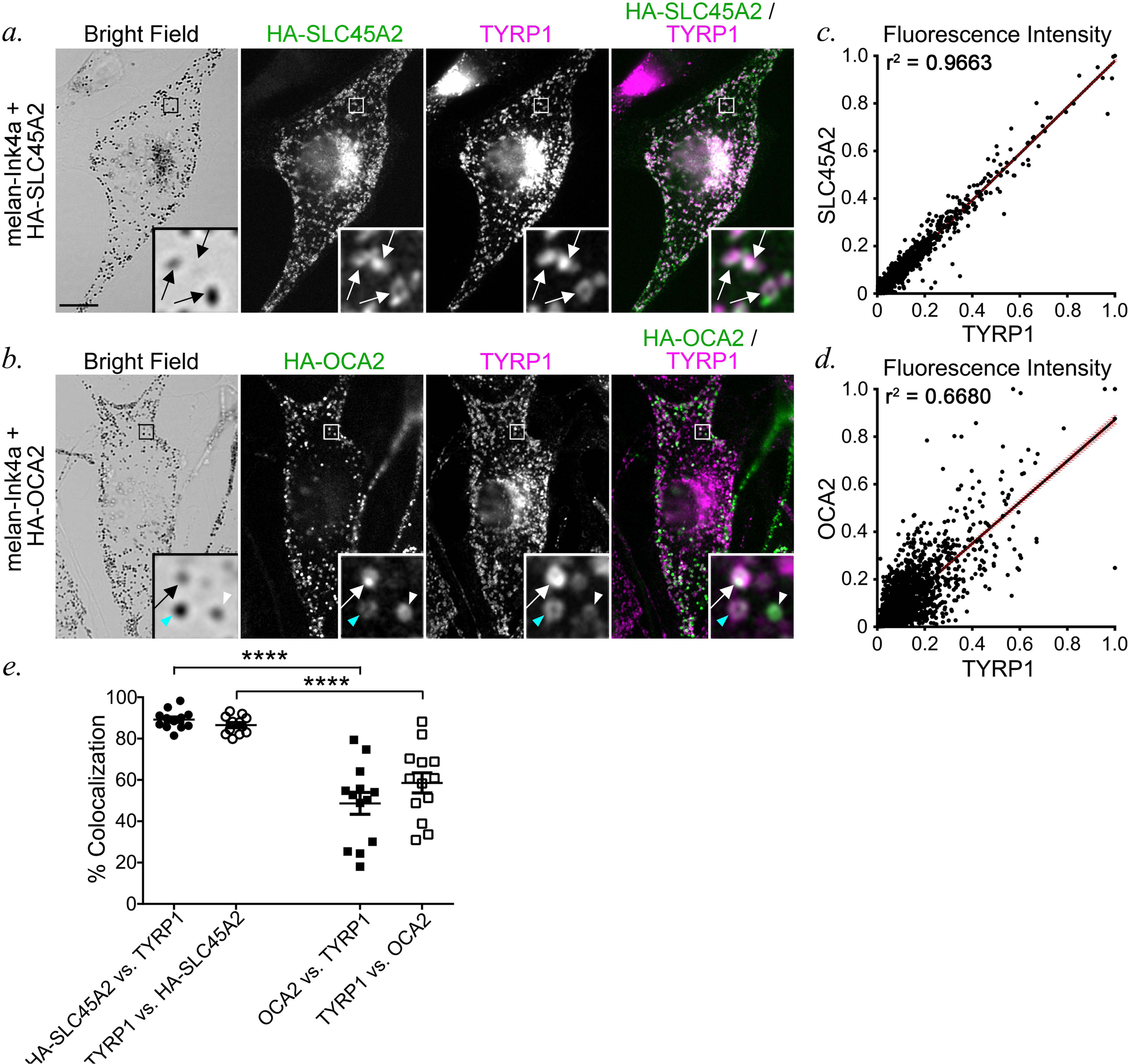
SLC45A2 and OCA2 occupy partially distinct melanosome subsets. **(*a*,** *b*) melan-Ink4a cells transiently expressing either HA-SLC45A2 (*a*) or HA-OCA2 (*b*) were fixed, labelled for HA (green) and TYRP1 (magenta), and analyzed by dIFM and bright field microscopy. Insets of boxed regions are magnified 7.5 times. Scale bar, 10 μm. White arrows, colocalization of HA-SLC45A2 or HA-OCA2 to TYRP1-containing compartments at comparable relative intensities; white arrowhead, compartments with high HA-OCA2 and low TYRP1; cyan arrowhead, compartments with low HA-OCA2 and high TYRP1. (*c*, *d*) Quantification of the object-based fluorescence intensity of HA-SLC45A2 versus TYRP1 (*c*) or HA-OCA2 versus TYRP1 (*d*) in at least 10 cells from 2 independent experiments. Correlation coefficients (r^2^) of fluorescent intensities for each pair of markers is indicated. (*e*) Manual quantification (mean ± SEM) of the % colocalization between TYRP1 and either HA-SLC45A2 or HA-OCA2 and TYRP1, shown as a percentage of total TYRP1-, SLC45A2-, or OCA2-containing compartments. Values are presented as label 1 vs label 2, in which compartments containing both label 1 and label 2 is indicated as a percentage of total compartments containing label 2. Data are quantified from at least 12 cells from 2 independent experiments. Statistical significance was determined using one-way ANOVA with Sidak’s test for multiple comparisons; only significant differences are indicated. ****, p < 0.001.

The distinct localization patterns for OCA2 and SLC45A2 suggest that they might regulate the lumenal pH and pigmentation of melanosomes at different maturation stages. We therefore tested for potential differences in melanosome pigmentation in melan-uw cells and immortalized melan-p1 melanocytes from OCA2-deficient pink-eyed dilute mice. Electron microscopy analyses showed that relative to either WT melan-Ink4a or “rescued” melan-uw cells stably expressing HA-SLC45A2 (melan-uw:HA-SLC45A2), both melan-uw and melan-p1 cells harbored melanosomes of similar size and number, but with more stage III melanosomes and fewer stage IV melanosomes (**Figure 5a-e**). However, only melan-p1 cells contained significantly more stage I/II melanosomes than the controls (**Figure 5e**), and the amount of pigment in stage III melanosomes in melan-p1 cells was lower than in melan-uw cells (**Figure 5**, compare panels **b** and **c**). Consistently, while both melan-uw and melan-p1 had dramatically lower melanin content than melan-Ink4a or melan-uw:HA-SLC45A2 cells, the melanin content of melan-p1 cells was significantly lower than that of melan-uw (**Figure 5f**). Taken together, these data suggest that OCA2 is required at an earlier stage of melanization than SLC45A2 during melanosome maturation.

**Figure 5.**
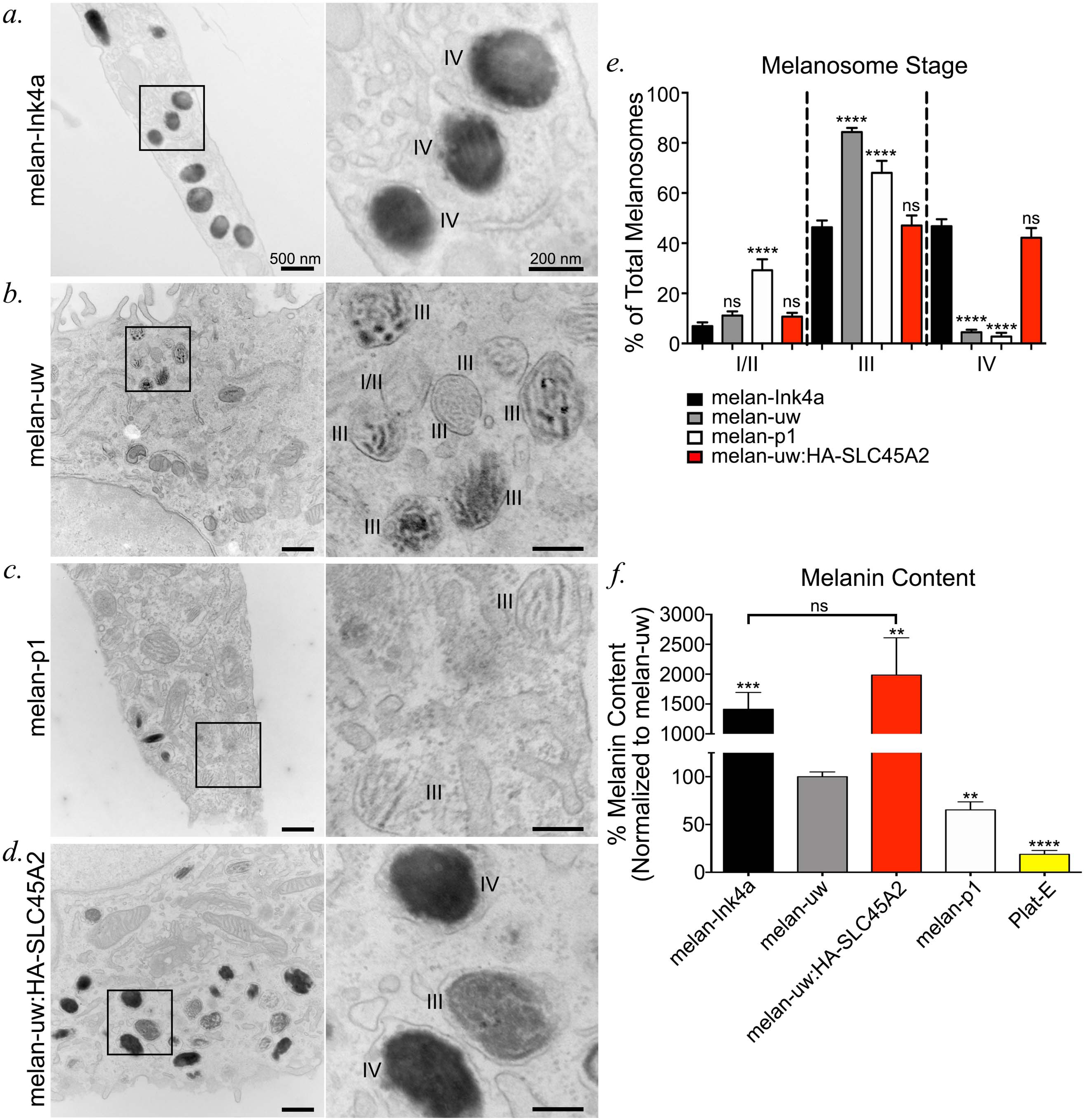
SLC45A2 functions at a later stage than OCA2 to facilitate melanosome maturation. (*a*-*d*) melan-Ink4a (*a*), melan-uw (*b*), melan-p1 (*c*), and melan-uw cells stably expressing HA-SLC45A2 (melan-uw:HA-SLC45A2; *d*) were fixed and analyzed by conventional transmission electron microscopy. Insets of boxed regions on the left were magnified 4.5 times on the right. I/II, stage I/II melanosomes; III, stage III melanosomes; IV, stage IV melanosomes. Scale bars, 500 nm (left) and 200 nm (right). (e) The number of stage I/II, stage III, and stage IV melanosomes per cell was quantified for each of the cell lines and is shown as a percentage of total counted melanosomes per cell. For each stage, data represent the mean ± SEM from at least two independent experiments where at least 20 cells and greater than 600 melanosomes were analyzed. Statistical analyses were performed by one-way ANOVA with multiple comparisons to melan-Ink4a for each stage and Sidak’s test for multiple comparisons. (*f*) Melanin content in cell lysates was determined by spectrometry relative to total protein content. The data are presented as a percentage normalized to melanin content in melan-uw samples and represent n>3 independent experiments, each performed at least in duplicate. Statistical significance was determined by Student’s two-sample t-test assuming unequal variances with FDR correction for multiple comparisons. **, p < 0.01; ***, p < 0.005; ****, p < 0.001; ns, no significant difference.

### OCA2 overexpression partially compensates for loss of SLC45A2 expression

Given that both OCA2 and SLC45A2 are capable of raising lysosomal pH when expressed in HeLa cells, we tested whether overexpression of either protein could compensate for the loss of the other. HA-SLC45A2, HA-OCA2, or HA-SLC35D3 as a negative control (see **Figure 1**) were transiently expressed by transfection in either *Slc45a2*-deficient melan-uw melanocytes or *Oca2*-deficient melan-p1 melanocytes, and the fraction of HA-positive cells that were pigmented after 3 days was quantified. As expected, very few melan-uw cells expressing HA-SLC35D3 were pigmented above background (19.88 ± 5.76%) whereas most HA-SLC45A2 expressing cells were pigmented (70.7 ± 5.85%; **Figure 6a, b, g**). Surprisingly, a large fraction of HA-OCA2-expressing melan-uw cells were also pigmented (46.37 ± 6.29%; **Figure 6c, g**), suggesting that OCA2 overexpression can compensate for the loss of SLC45A2. By contrast, whereas HA-OCA2 expression restored pigmentation to melan-p1 cells (93.17 ± 4.97%), HA-SLC45A2 expression was as ineffective at restoring pigmentation as the negative control HA-SLC35D3 (**Figure 6d-f, h**). Thus, overexpression of SLC45A2 cannot compensate for the loss of OCA2. These data are consistent with a model in which sequential function of OCA2 and then SLC45A2 is required to modulate and maintain a nearly-neutral pH in melanosomes.

**Figure 6.**
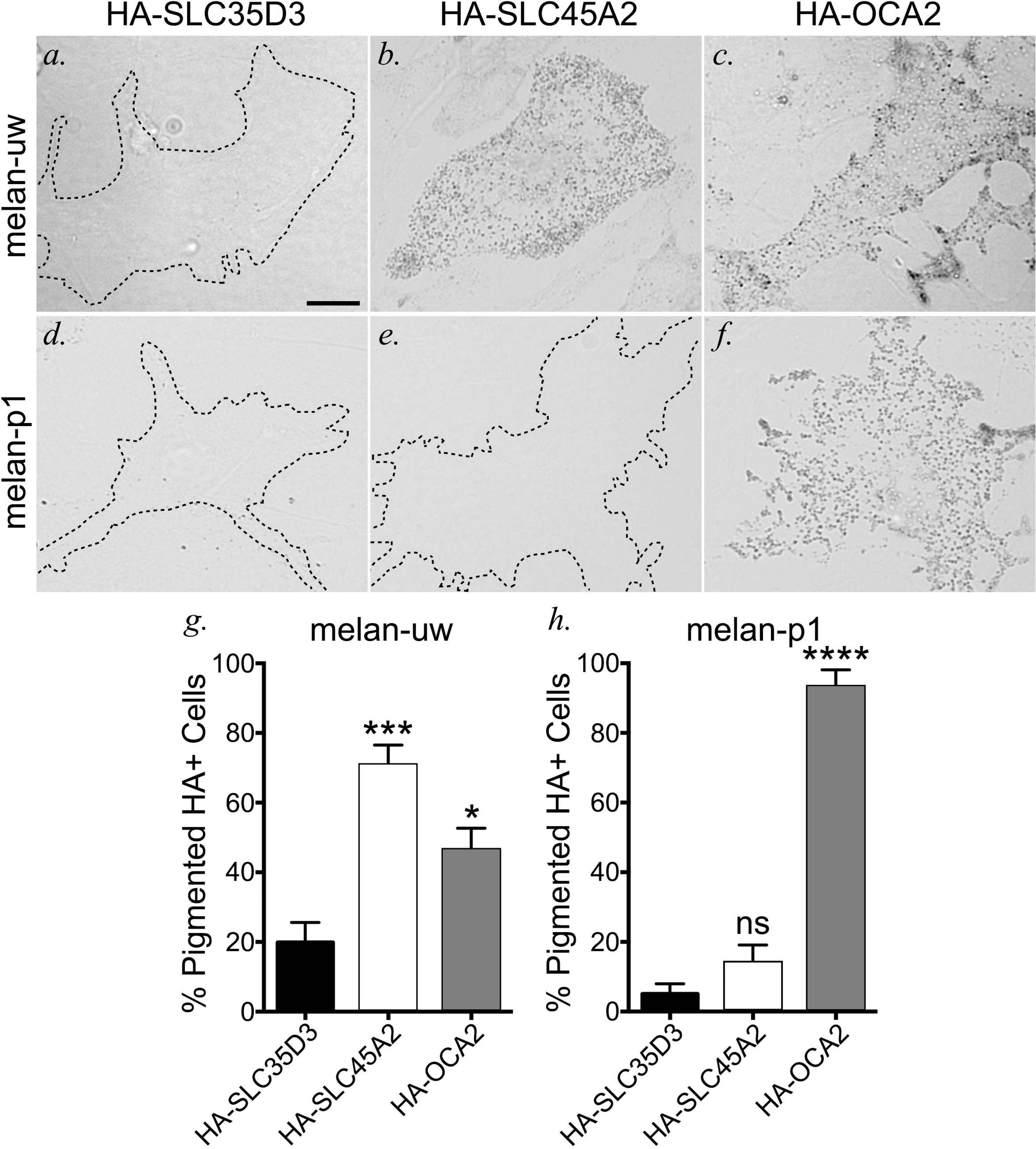
OCA2 overexpression compensates for loss of SLC45A2-dependent melanosome neutralization in melanocyte pigmentation. SLC45A2-deficient melan-uw cells or OCA2-deficient melan-p1 cells were transiently transfected with negative control HA-SLC35D3 (*a*, *d*), HA-SLC45A2 (*b*, *e*), or HA-OCA2 (*c*, *f*), and fixed and labelled for HA (not shown). Bright field microscopy was used to visualize pigmented melanosomes within HA-positive cells. Scale, 10 μm. (*g*, *h*) The number of HA-containing pigmented cells was quantified and is depicted as a mean percentage of total HA-containing (HA+) cells ± SEM. Quantification is from at least 3 independent experiments. Statistical significance was determined using one-way ANOVA with Holm-Sidak’s test for multiple comparisons. *, p < 0.05; ***, p < 0.005; ****, p < 0.001; ns, not significant.

### The light skin-associated F374 SLC45A2 variant protein is expressed at lower levels than the dark skin-associated L374 variant

The major human *SLC45A2* allele associated with light skin tone encodes a phenylalanine (F) instead of a leucine (L) at amino acid position 374, within the eighth transmembrane domain of the predicted 12-transmembrane domain-containing protein (**Figure 7a**). How does this single amino acid substitution impact pigmentation? We reasoned that the F374 variant could affect either protein folding/stability, localization to melanosomes, or transporter function within melanosomes. To begin to distinguish between these possibilities, we generated stable melan-uw cells expressing the HA-tagged dark skin-associated L374 variant (as used in **Figures 1-6**), the HA-tagged F374 variant, or HA-SLC35D3 as a control (CTRL). Bright field microscopy analysis revealed that cells expressing HA-SLC45A2-F374 were darker than control cells but much lighter than cells expressing HA-SLC45A2-L374 (**Figure 7b**), consistent with published results (Cook, Chen et al., 2009). The observed phenotype was confirmed by quantitative melanin content assay, in which cells expressing HA-SLC45A2-F374 harbored significantly more pigment than control cells but less than cells expressing HA-SLC45A2-L374 (**Figure 7c**). The lower level of pigmentation in cells expressing HA-SLC45A2-F374 did not reflect reduced SLC45A2 mRNA expression, as quantitative RT-PCR analyses showed that melan-uw cells expressing HA-SLC45A2-F374 had 1.25-fold more SLC45A2 mRNA than the wild-type L374-expressing cells (**Figure 7d**). However, immunoblotting analyses of cell lysates for the HA-tagged proteins revealed that the L374 stable cell line expressed >4-fold more HA-SLC45A2 than cells stably expressing the F374 variant (**Figure 7e, f**). These data indicate that the reduced pigmentation conferred by F374 is reflected by a post-translational reduction in SLC45A2 protein content.

**Figure 7.**
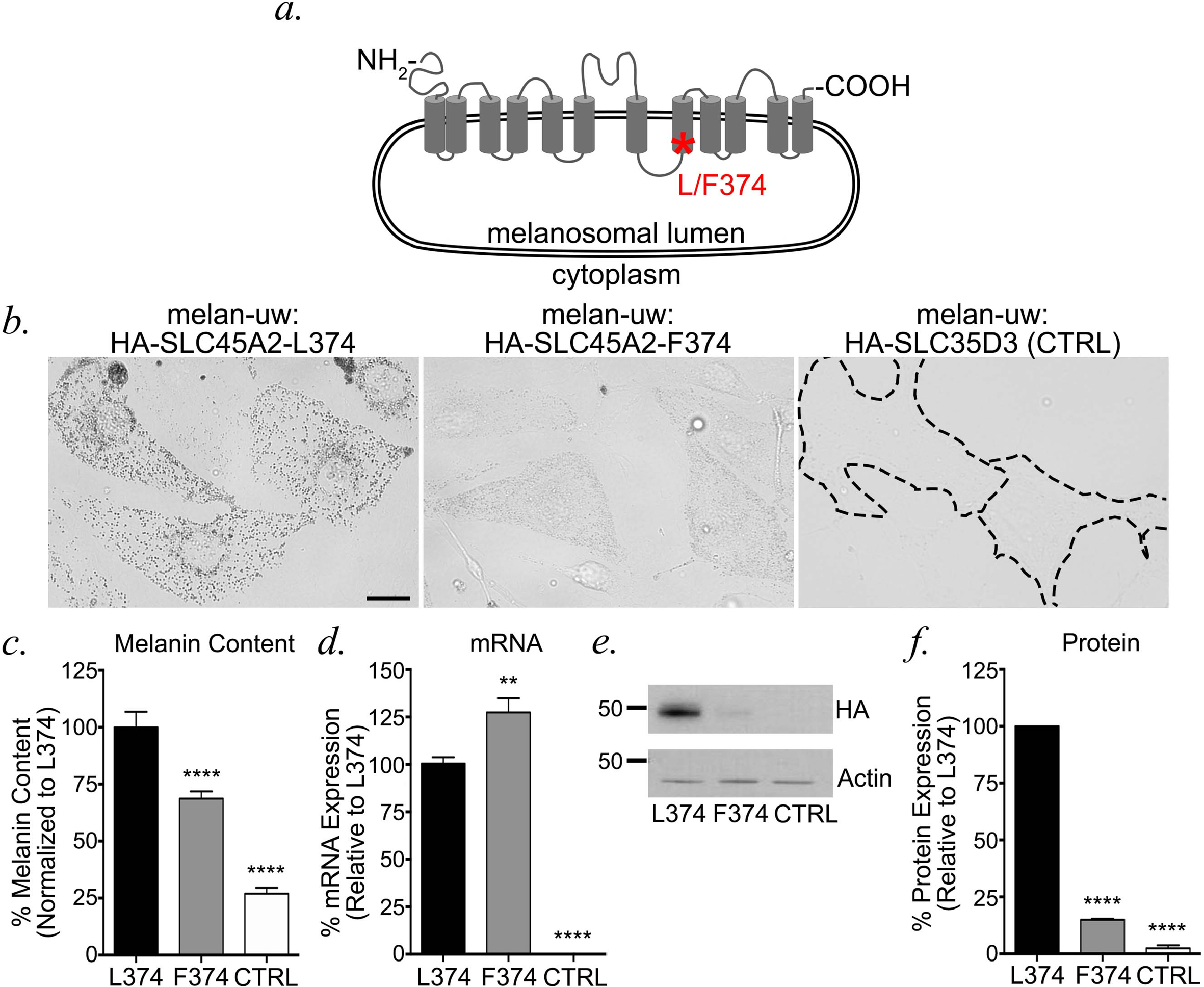
The human light skin-associated SLC45A2-F374 variant is expressed in melanosomes at lower levels than the dark variant. (*a*) Schematic diagram of human SLC45A2 primary structure, with transmembrane (gray cylinders), luminal and cytosolic domains (gray lines) indicated. An asterisk indicates the position of L/F374. (*b*) Stably transfected melan-uw cells expressing HA-SLC45A2-L374 (left), HA-SLC45A2-F374 (center), or HA-SLC35D3 as a negative control (right) were imaged by bright field microscopy to emphasize pigmentation levels. Bar, 10 μm. (*c-f*) Stably transfected melan-uw cells expressing HA-SLC45A2-L374 or -F374 or HA-SLC35D3 (CTRL) were analyzed for (*c*) melanin content by quantitative spectroscopy assay, (*d*) SLC45A2 mRNA expression by quantitative RT-PCR, or (*e*, *f*) HA-SLC45A2 protein expression by immunoblotting relative to actin as a loading control. A representative immunoblot is shown in *e*; quantification of SLC45A2 expression normalized to actin levels over three experiments is shown in *f*. All values are normalized to 100% for cells expressing HA-SLC45A2-L374. Each experiment was repeated three times. Statistical analyses were determined by one-way ANOVA with Dunnett’s test for multiple comparisons. **, p < 0.01; ****, p < 0.001.

To test whether the reduced protein content of the F374 variant reflected increased protein degradation, we assessed the protein stability of each variant over time in stably transduced melan-uw cells. Cells were treated with cycloheximide (CHX) to block new protein synthesis, and the amount of HA-SLC45A2 remaining in cell lysates at different time points was assessed by immunoblotting. While the levels of both variants were substantially reduced by 32 h following CHX treatment, HA-SLC45A2-F374 was degraded at a higher rate than HA-SLC45A2-L374 (L374 half-life, 16.5 ± 3.5 h; F374 half-life, 8.9 ± 0.2 h) and was nearly completely eliminated by 32 h (**Figure 8a, c**). To determine whether the rapid decline in the F374 variant protein levels required proteasomal degradation, cells were treated with the proteasome inhibitor MG132 during the CHX chase. Surprisingly, MG132 nearly completely blocked the gradual disappearance of HA-SLC45A2-L374, but did not impede the rapid degradation of HA-SLC45A2-F374 at early chase times (**Figure 8b, c**). MG132 treatment did, however, block the slow degradation of F374 at later time points (**Figure 8b, c**). These data indicate that normal turnover of L374 and of the stable fraction of F374 requires proteasome activity, but that rapid degradation of F374 occurs by a proteasome-independent mechanism. Importantly, dIFM analysis showed that HA-SLC45A2-F374 colocalized with TYRP1 to a similar extent as HA-SLC45A2-L374 (**Figure 8d, e**), indicating that the cohort of non-degraded F374 variant localizes properly to melanosomes where it retains some function in supporting elevated pH.

**Figure 8.**
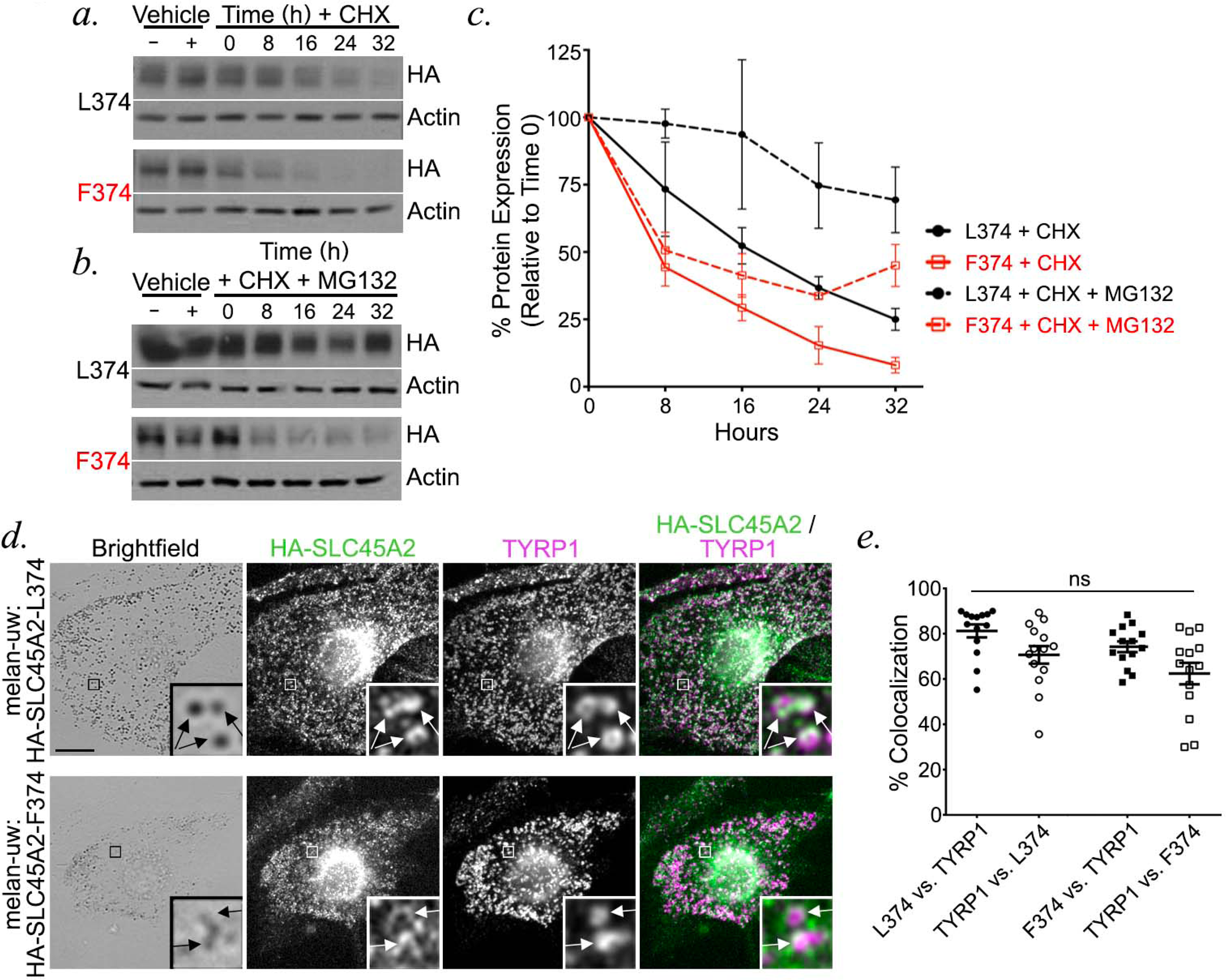
The F374 variant is less stable than the L374 variant and is degraded by a proteasome-independent pathway. (*a, b*) Melan-uw cells stably expressing either HA-SLC45A2-L374 (L374; *a*, *b*, top) or -F374 (F374; *a, b,* bottom) were cultured with cycloheximide (CHX) alone (*a*) or combined with proteasome inhibitor MG132 (*b*) for the indicated times (hours, h). Control cells were incubated for 32 h with vehicle controls (EtOH for CHX; DMSO for MG132). At the indicated times, cells were harvested and whole cell lysates were fractionated by SDS-PAGE and analyzed by immunoblotting for HA and for actin as a loading control. Relevant regions of the gels are shown. (*c*) Band intensities for HA-SLC45A2 were quantified over three independent experiments, normalized to actin band intensities, and presented as a percentage of the signal observed at time 0 +/-SEM. (*d, e*) Melan-uw cells stably expressing either HA-SLC45A2-L374 (*d*, top) or HA-SLC45A2-F374 (*d,* bottom) were fixed, labeled for HA (green) and TYRP1 (magenta), and analyzed by dIFM and by bright field microscopy. Merged HA/TYRP1 image is at right; insets of boxed regions are magnified 7.5 times. HA-SLC45A2 and TYRP1 localizing to the same compartments are indicated with arrows. Scale, 10 μm. (*e*) The degree of TYRP1 and SLC45A2 localization to the same compartments was quantified manually from 14 cells over 2 independent experiments, and is presented as label 1 vs label 2, in which the colocalization between label 1 and label 2 is a percentage of total label 2. Statistical significance was determined using one-way ANOVA with Sidak’s test for multiple comparisons. ns, no significant difference.

Taken together, these data suggest that the light skin-associated F374 variant of SLC45A2 is less stable than the dark skin-associated L374 variant due to non-proteasomal degradation, resulting in lower SLC45A2 activity in melanosomes.

## DISCUSSION

SLC45A2 is a critical determinant of skin and eye pigmentation, but until now its localization and functional role in melanogenesis have not been clearly delineated. Using a functional epitope-tagged form of SLC45A2 expressed in immortalized SLC45A2-deficient epidermal mouse melanocytes, we show here that SLC45A2 localizes to a subset of mature melanosomes marked by TYR and TYRP1 expression, where it partially accumulates in subdomains of the organelle outer membrane. When expressed ectopically in HeLa cells, SLC45A2 localizes to subdomains of the lysosomal membrane and functions to increase lysosomal pH, supporting a previously proposed role in neutralizing melanosomes (Bin et al., 2015). Although the melanosomal chloride channel OCA2 also functions to neutralize melanosome pH (Bellono et al., 2014), SLC45A2 localizes to a partially distinct cohort of melanosomes, and SLC45A2-deficient melanocytes harbor melanosomes that are more pigmented than OCA2-deficient cells – suggesting that SLC45A2 functions at a later melanosome maturation stage than OCA2. Accordingly, overexpression of OCA2 in SLC45A2-deficient melanocytes compensated for SLC45A2 deficiency in pigment production, whereas SLC45A2 overexpression could not compensate for OCA2 deficiency. Taken together, these functional data lead us to conclude that SLC45A2 functions to neutralize lumenal pH at a later stage of melanosome maturation than OCA2. We also identified the mechanism underlying the pigmentation phenotype of the light skin-associated F374 variant of SLC45A2. Although it localizes to melanosomes like the dark skin-associated L374 variant, SLC45A2-F374 is less stable and more rapidly degraded than -L374, resulting in markedly reduced SLC45A2-F374 protein levels at steady state. Together, our data (1) indicate that SLC45A2 regulates melanogenesis by controlling melanosome pH in a manner that is non-redundant with OCA2 and (2) uncover the mechanism by which a common genetic variant imparts light skin color.

Our data indicate that SLC45A2 localizes to the melanosomal membrane, where we propose that it directly functions to increase lumenal pH. This is analogous to other proteins on the melanosome membrane, such as the chloride channel OCA2 and the cation channel TPC2, which also directly regulate melanosome pH through ion transport (Ambrosio et al., 2016, Bellono et al., 2014, Bellono et al., 2016, Sitaram et al., 2009), and contrasts with ion transporters/channels that may indirectly regulate melanosome pH through effects on other organelles. For example, the putative solute transporter MFSD12 negatively regulates pigmentation from lysosomes (Crawford et al., 2017) and NCKX5 encoded by *SLC24A5*, the gene deficient in OCA6, appears to positively regulate pigmentation from the trans Golgi network (Ginger, Askew et al., 2008, Rogasevskaia, Szerencsei et al., 2019). Together, these observations suggest that melanosome pH is regulated both directly and indirectly via a complex network of putative channels/transporters. Further work is necessary to identify other regulators of melanosome pH and to understand how these proteins cooperate to fine-tune the lumenal pH of melanosomes, thereby controlling melanin synthesis.

When expressed at the plasma membrane in yeast, SLC45A2 functions as a sucrose/proton symporter that transports sucrose from the extracellular space into the cytosol in a manner that requires a proton gradient and is maximal at a slightly acidic pH (Bartölke et al., 2014). Interestingly, SLC45A2 and other SLC45 family members can transport not only the disaccharide sucrose but also the monosaccharides glucose, fructose, and mannose (Bartölke et al., 2014). If it were to function similarly in melanocytes, SLC45A2 would expel protons and sugar molecules from the melanosome lumen into the cytosol, hence increasing melanosome pH. Whether sugars are substrates for SLC45A2 under physiological conditions in melanosomes remains to be determined, but given that lysosomal enzymes are present within maturing melanosomes (Diment, Eidelman et al., 1995, Raposo et al., 2001), it is possible that monosaccharides released from glycoproteins by lysosomal glycosidases in early stage melanosomes could serve as substrates for sugar/proton symporter activity. Our data show that expression of SLC45A2 in lysosomes of HeLa cells is sufficient to increase pH, indicating that its proton-dependent symporter activity is sufficient to raise lumenal pH by counteracting the activity of the vacuolar ATPase (vATPase), which pumps protons into the lumen of both melanosomes and lysosomes in an ATP-dependent manner (Bhatnagar & Ramalah, 1998, Mindell, 2012, Tabata, Kawamura et al., 2008). Sugar co-transport from the melanosome into the cytosol would also decrease the osmotic concentration inside the melanosome, requiring counter-ion transport to balance solute concentrations. Further investigation into the melanosomal channels, transporters, and exchangers is required to understand how the melanosome regulates not only its pH but also its osmolarity.

Unlike melanosomal proteins such as TYRP1 that are detected relatively uniformly on the melanosome membrane by IFM, SLC45A2 is sometimes detected in a punctate pattern, corresponding to specific subdomains of the melanosome membrane. The nature of these subdomains and the mechanism by which SLC45A2 assembles into them remains unclear. The punctate localization is conserved on lysosomes upon ectopic expression of SLC45A2 in non-pigmented HeLa cells, suggesting that the assembly of SLC45A2 into subdomains does not require interactions with melanocyte-specific proteins. Some cells expressing HA-SLC45A2 displayed a mixture of punctate and ring-like SLC45A2 patterns on melanosomes. This might either reflect an artifact of overexpression or the consequence of a regulated process of aggregate assembly that may be influenced by melanosome contents or other features of melanosome maturation. The composition, regulation and function of these SLC45A2-containing structures requires further investigation.

The high colocalization between SLC45A2 and TYRP1 suggests that SLC45A2 is co-delivered with TYRP1 to maturing melanosomes, likely via BLOC-1-dependent membrane transport (Setty, Tenza et al., 2008, Setty, Tenza et al., 2007). In contrast, there is significantly less overlap between OCA2 and TYRP1, despite similar requirements for BLOC-1 for delivery to melanosomes (Sitaram et al., 2012), and their abundance is inversely related in structures in which they overlap. Moreover, the relative melanin content of SLC45A2- and OCA2-deficient melanocytes is different, such that SLC45A2-deficient melanocytes harbor more mature melanosomes and slightly higher pigmentation than OCA2-deficient melanocytes. These results suggest that OCA2 functions at an earlier melanosomal maturation stage compared to SLC45A2, implying either that the melanosome delivery of OCA2 and TYRP1/SLC45A2 is staggered or that OCA2 is cleared soon after delivery. The latter is supported by the short half-life of OCA2 (Sitaram et al., 2009), the dependence of this half-life on melanization (Donatien & Orlow, 1995), and the accumulation of OCA2 on internal membranes in mature melanosomes (Sitaram et al., 2012). Based on these observations, we propose a model in which OCA2, SLC45A2, and TYRP1 are delivered together to maturing melanosomes - which harbor active vATPase to acidify their lumen (Bhatnagar & Ramalah, 1998, Tabata et al., 2008). OCA2 would facilitate a rapid efflux of chloride ions, immediately reducing membrane potential and thereby slowing vATPase activity. As the melanosomes mature, OCA2 clearance from the limiting membrane by invagination onto internal membranes coupled with continued SLC45A2 arrival would support a slow but continued melanosome neutralization mediated by SLC45A2 proton export, counteracting residual vATPase activity. If this model were true, then OCA2 overexpression, by prolonging exposure on the melanosomal membrane, would antagonize vATPase activity in a more sustained way, perhaps resulting in slow proton efflux from the melanosome through another means such as membrane recycling (Dennis, Delevoye et al., 2016). This model would thus explain our observation that OCA2 overexpression can rescue SLC45A2 loss but that SLC45A2 overexpression - with its limited capacity for proton export - cannot rescue OCA2 loss.

The SLC45A2-F374 variant is the predominant allele in Northern Europe and is strongly associated with light skin and hair and blue eyes (Branicki et al., 2008, Cook et al., 2009, Lao, de Gruijter et al., 2007, Lopez et al., 2014, Lucotte, Mercier et al., 2010, Lucotte & Yuasa, 2013, Mukherjee, Mukerjee et al., 2013, SabetiVarilly et al., 2007, Soejima & Koda, 2007, Soejima, Tachida et al., 2006, Yuasa et al., 2006). However, the mechanism responsible for the reduced pigmentation associated with SLC45A2-F374 was previously unclear. We show here that compared to the SLC45A2-L374 allele, expression of the F374 allele in SLC45A2-deficient melanocytes results in a much more meager increase in pigmentation, despite higher levels of *SLC45A2* mRNA. Our results demonstrate that this is due to instability of the SLC45A2-F374 protein, leading to a higher rate of degradation relative to SLC45A2-L374 through a proteasome-independent mechanism. Nevertheless, the remaining cohort of SLC45A2-F374 localizes appropriately to melanosomes. Thus, we propose that the reduced pigmentation associated with the F374 allele is due to insufficient levels of SLC45A2-F374 on the melanosomal membrane to maintain optimal neutral pH for maximal TYR activity, thus resulting in decreased melanin synthesis. Whether the F374 variant is rapidly degraded following melanosomal delivery or is degraded in a distinct compartment remains to be determined.

In conclusion, our data significantly advance our understanding of SLC45A2 and its function on melanosomes and begin to unravel the stepwise contributions of SLC45A2 and OCA2 to lumenal pH regulation and proper melanosome maturation. We also reveal the molecular basis explaining why the SLC45A2-F374 variant is a hypomorphic allele and propose a mechanism by which this allele contributes to lighter pigmentation in humans. The regulation of melanosome pH through the coordinated function of transporters, channels, and other proteins requires further investigation.

## MATERIALS AND METHODS

### Reagents

Unless otherwise specified, chemicals were obtained from Sigma-Aldrich (St. Louis, MO) and tissue culture reagents from Life Technologies/ Thermo Fisher Scientific (Waltham, MA). Protease inhibitors were purchased from Roche Diagnostics (Rotkreuz, Switzerland), gene amplification primers from Integrated DNA Technologies (Coralville, IA), GoTaq DNA polymerase from Promega Corp. (Madison, WI), and restriction enzymes and T4 DNA ligase from New England Biolabs (Ipswich, MA).

### Cell culture

Immortalized melanocyte cell lines melan-p1 (OCA2 deficient) from C3H-*Oca2^cp^*/*^25H^* (pink-eyed dilute) mice (Sviderskaya, Bennett et al., 1997), and “wild-type” (WT) melan-Ink4a-Arf1 (formerly called melan-Ink4a-1; referred to here as melan-Ink4a or WT) from C57BL/6J-*Ink4a-Arf^−/−^* (*Cdkn2* null) mice (Sviderskaya et al., 2002) have been described and were derived from the skins of neonatal mice. Three immortal SLC45A2 deficient melanocyte lines, melan-uw-1, −2 and −3, were similarly derived from the skins of neonatal C57BL/6J-*Ink4a-Arf^-/-^ Slc45a2^uw/uw^* (underwhite) mice. Mice were housed and interbred by Dr Lynn Lamoreux (Texas A&M University, College Station, TX, USA), and skins were shipped on ice to London. Only the melan-uw-2 line (referred to here as melan-uw) was used here because they were more uniform in pigmentation both before and after stable expression of HA-SLC45A2. All cells were cultured at 37°C and 10% CO_2_ in RPMI 1640 medium supplemented with 10% FBS (Atlanta Biologicals) and 200 nM 12-*O*-tetradecanoylphorbol-13-acetate (TPA). Melan-uw and melan-p1 cells were additionally supplemented with 200 pM cholera toxin.

Retrovirus production from transiently transfected Plat-E cells(Morita, Kojima et al., 2000) and retroviral transduction of melanocyte cell lines were carried out as described previously (Meng, Wang et al., 2012, Setty et al., 2007). Briefly, Plat-E cells were transfected with retroviral DNA constructs using Lipofectamine 2000 (Thermo Fisher), and the medium was replaced the next day. Retrovirus-containing supernatants were collected 48 h later, filtered, and added to melan-Ink4a cells or melan-uw cells in a 1:1 ratio with fresh medium. The medium was replaced the next day, and pools of stable transductants were selected 24 h later by adding 300 μg/ml hygromycin B to the medium. Stable transfectants were occasionally treated with 200 µg/ml hygromycin B for 2–3 d to maintain selective pressure for the transgene. For transient transfections, cells on glass coverslips were transfected with DNA constructs using Lipofectamine 3000 or Lipofectamine 2000 (Thermo Fisher), and the medium was replaced the next day. Cells were fixed using 4% formaldehyde/PBS 48 or 72 hours after transfection and stored at 4°C for immunolabeling and analysis.

### DNA Constructs

We first generated a GFP-SLC45A2 (L374) fusion protein by inserting human SLC45A2 cDNA, generated by RT-PCR of RNA isolated from darkly pigmented human epidermal melanocytes, into the BamHI/Xhol sites of pcDNA4/TO (Invitrogen/Life Technologies) in frame with GFP and a Gly-Ala-Gly-Ala linker previously inserted in the AflII/HindIII sites (sequence available upon request). To generate HA-tagged SLC45A2, the insert from pcDNA4/TO-GFP-SLC45A2 was amplified by PCR adding XhoI and NotI restriction sites at the 5’ and 3’ ends, respectively, and subcloned into the respective sites of pCI (Promega). An N-terminal Kozak consensus start site, HA tag, and Gly-Ser linker were subsequently added by PCR using a distinct forward primer (5’-gcatatctcgagatgTACCCATACGATGTTCCAGATTACGCTggctcagg atctgggatgggtagcaacagtgggc-3’) and the same reverse primer. The HA-SLC45A2 insert was subsequently subcloned as a XhoI-NotI fragment into pBMN-(XN)-IRES-hygro to generate the retroviral vector encoding HA-SLC45A2. The F374 variant was made by site-directed mutagenesis of the pBMN-(XN)-IRES-hygro-HA-SLC45A2 (L374) construct using the Clontech site-directed mutagenesis kit.

The plasmid-based expression vector pCR3-OCA2-WT-HA-UTR2 and corresponding retroviral vector pBMN-OCA2-WT-HA-UTR2 encoding human OCA2 with an exofacial HA epitope were described in (Sitaram et al., 2009); pCDM8.1-HA-SLC35D3 encoding human SLC35D3 with an N-terminal HA epitope tag was described in (Meng et al., 2012); and human LAMP1-mCherry in pCDNA3.1 was a generous gift from Sergio Grinstein (Univ. of Toronto and Hospital for Sick Children, Toronto, ON, Canada).

### Melanin Content Quantification

Melanin quantification by spectroscopy was done essentially as described (Delevoye, Hurbain et al., 2009). Briefly, melanocytes seeded on 6-cm dishes were trypsinized, pelleted, and sonicated in melanin buffer (50 mM Tris, 2 mM EDTA, and 150 mM NaCl, pH 7.4) supplemented with protease inhibitor cocktail (Roche). Insoluble material was pelleted for 15 min at 16,000 g (4°C), rinsed in ethanol/diethyl ether (1:1), and dissolved in 2 M NaOH/20% DMSO at 60°C. The optical density at 492 nm was measured to estimate melanin content and normalized to protein concentration as determined by BCA protein determination kit (Thermo Fisher). Plat-E cells, an unpigmented cell line, were used as a negative control. Statistical significance from at least 3 independent experiments was determined either by Student’s two-sample t-test assuming unequal variances with FDR correction for multiple comparisons or by one-way ANOVA with Dunnett’s test for multiple comparisons.

### Antibodies

Primary antibodies used and their sources (listed in parentheses) include: mouse monoclonal antibody TA99/Mel5 to TYRP1 (American Type Culture Collection; Rockville, MD); rat monoclonal antibody 3F10 to the HA11 epitope (Sigma); mouse monoclonal antibody H4A3 to human LAMP1 (Developmental Studies Hybridoma Bank, Iowa City, IA); rabbit anti-LAMP2 (Abcam; for mouse melanocytes); rabbit anti-TYR (Pep7h, to the C-terminal 17 amino acids of human TYR (Calvo et al., 1999)); mouse anti-beta actin (BA3R) monoclonal antibody (ThermoFisher Scientific). Species- and/or mouse isotype–specific secondary antibodies from donkey or goat and conjugated to Alexa Fluor 488, Alexa Fluor 594 used in IFM or conjugated with HRP for western blots were obtained from Jackson ImmunoResearch Laboratories (West Grove, PA).

### Bright field microscopy, immunofluorescence microscopy (IFM), and colocalization analyses

IFM analyses of fixed cells were done essentially as described (Dennis et al., 2016). Briefly, cells were plated on Matrigel (BD)-coated coverslips, fixed with 4% formaldehyde (VWR) in PBS, labeled with primary and secondary antibodies diluted in PBS/ 0.02% saponin/ 0.01% BSA, mounted onto slides using Prolong Gold (ThermoFisher), and analyzed on a Leica DMI-6000 microscope equipped with a 40× or 63× objective lens (Leica; 1.4 NA), a Hamamatsu Photonics ORCA-Flash4.0 sCMOS C11440-22CU digital camera, and Leica Application Suite X (LAS X) software. Images in sequential z planes (0.2 μm step size) were deconvolved with Microvolution software and further analyzed using ImageJ (http://fiji.sc/Fiji; National Institutes of Health).

Because SLC45A2 localized partially to subdomains on the melanosome membrane and thus did not fully overlap other melanosomal markers, previously used automated methods to quantify colocalization did not yield meaningful values; similarly, OCA2 localized largely to the interior of melanosomes(Sitaram et al., 2012, Sitaram et al., 2009) and was thus partially distinct from the distribution of TYRP1 or TYR cytoplasmic domain on the melanosome membrane. Therefore, quantification of the degree of overlap between two channels on individual organelles was performed manually on deconvolved, Z-projected images of cells in ImageJ. In brief, single-channel images and a merged image were synchronized using “Analyze > Tools > Synchronize Windows;” signals that were localized to the same organelle were counted using the “Multi-Point” tool; remaining signal in single channels were counted using the “Multi-Point” tool; and the total colocalized structures was calculated as a percentage of total structures for each channel. Statistical significance was determined using one-way ANOVA with Sidak’s multiple comparisons test.

Scoring of pigmentation in transfected melan-uw and melan-p1 cells was done blinded. Fixed, stained coverslips were randomly assigned numbers and samples were revealed only after images were taken and scored. Statistical significance was determined using one-way ANOVA with Holm-Sidak’s test for multiple comparisons.

### Fluorescence intensity analyses

Object-based fluorescence intensity was measured using ImageJ as described in https://www.unige.ch/medecine/bioimaging/files/1914/1208/6000/Quantification.pdf. Briefly, 8-bit, single-channel images were opened in ImageJ and brightness and contrast adjusted for each image. The image was duplicated and a threshold applied using “Image > Adjust > Auto Local Threshold.” To multiply binary images of the two channels of interest, we used “Process > Image Calculator > Multiply.” To redirect intensity measurements from binary image to fluorescent image, we used “Analyze > Set measurements” and pulled down the “Redirect to:” menu to choose the appropriate fluorescent image, making sure “Area” and “Integrated density” were also checked. With “binary image” clicked, we used “Analyze > Analyze Particles” to measure object intensity. This was repeated for each channel, and for each sample values were normalized to the highest integrated density value in each channel. At least 10 cells from each of two independent experiments were analyzed.

### Lysosomal pH measurements

HeLa cells were transfected one day prior to imaging experiments with LAMP1-mCherry alone, mCherry-OCA2 alone, or HA-SLC45A2 and LAMP1-mCherry to identify endolysosomes, and incubated with 1 μM LysoSensor-ND160 (Thermo Fisher) for 5 min. Lysosensor was excited at 405 nm and its emission detected at 417-483 nm (W1) and 490-540 nm (W2). The ratio of LysoSensor W1/W2 emission in endolysosomes expressing LAMP1-mCherry was assigned a pH value based on a calibration curve generated prior to each experiment using solutions containing 125 mM KCl, 25 Mm NaCl, 24 μM Monensin, and varying concentrations of MES to adjust the pH to 3.5, 4.5, 5, 5.5, 6.5, 7, 7.5. The fluorescence ratio was linear from pH 5 - 7.0. Statistical significance was determined by one-way ANOVA with Sidak’s test for multiple comparisons.

### Electron microscopy

Melanocytes were cultured in 10-cm dishes and fixed in situ with Karnovsky’s fixative [4% paraformaldehyde (VWR Scientific, Radnor, PA), 4 mM calcium chloride, 72 mM sodium cacodylate, pH 7.4] containing 0.5% glutaraldehyde (Polysciences, Warrington, PA) for 1–2 h at room temperature. This solution was then removed and replaced with Karnovsky’s fixative containing 2% glutaraldehyde, and the cells were fixed overnight at room temperature. Cells were collected by scraping using a cell scraper and centrifugation at 27 g for 5 min, resuspended in Karnovsky’s fixative containing 0.5% glutaraldehyde, and stored at 4°C until processing. After subsequent buffer washes, the samples were post-fixed in 2.0% osmium tetroxide with 1.5% K_3_Fe(CN)_6_ for 1 hour at room temperature, and then rinsed in deonized water prior to en bloc staining with 2% uranyl acetate. After dehydration through a graded ethanol series, the tissue was infiltrated and embedded in EMbed-812 (Electron Microscopy Sciences, Fort Washington, PA)) at the Electron Microscopy Resource Laboratory (University of Pennsylvania). Thin sections were stained with uranyl acetate and SATO lead and examined with a JEOL 1010 electron microscope fitted with a Hamamatsu digital camera (Hamamatsu, Bridgewater, NJ) and AMT Advantage NanoSprint500 software (Advanced Microscopy Techniques, Woburn, MA).

For melanosome staging, a combined total of at least 20 cells from at least two independent experiments and over 600 melanosomes were analyzed for each cell type. The number of stage I/II, stage III, and stage IV melanosomes were quantified, and the percentage of melanosomes at each stage was calculated for each cell. These data were analyzed and plotted using Prism 6 (GraphPad, La Jolla, CA), and statistical significance was determined by one-way ANOVA with multiple comparisons (samples compared to melan-Ink4a) and Sidak’s test for multiple comparisons.

### Immunoblotting

Melan-uw cells stably expressing HA-SLC45A2-L374, HA-SLC45A2-F374, or HA-SLC35D3 cultured in 60 mm Petri dishes were treated with 100 μg/ml cycloheximide and collected for western blot every 8 h for 32 h. For each time point the cells were rinsed in cold PBS before adding 40 μl RIPA buffer (EMD Millipore) with protease inhibitor mixture (Roche). Cells were scraped from Petri dishes, then homogenized using a 22G syringe. Lysates were rotated end-over-end for 1 h at 4°C, then centrifuged at 14,000 rpm for 30 min at 4°C to remove cell debris. The supernatant from each sample was kept at 4°C until all the time points were collected. Protein concentration for each sample was measured using the BCA assay (Pierce) according to the manufacturer’s protocol. 5 μg protein from each sample was fractionated on a 4-12% Bis-Tris SDS-PAGE gel (Life Technologies/ Thermo Fisher Scientific), then electrotransferred to nitrocellulose, blocked with 5% milk in PBS/ 0.5% Tween and incubated with antibodies to HA or β-actin diluted in PBS/ 0.5% Tween. The blots were washed with PBS/ 0.5% Tween and then the primary antibodies were detected by incubation with goat α-rat or α-mouse secondary antibodies coupled to HRP diluted in PBS/ 0.5% Tween. Statistical significance was determined using one-way ANOVA with Dunnett’s test for multiple comparisons. Protein half-lives ± SEM were calculated using interpolation with second order polynomials for each replicate, which provided curves that best matched the experimental data.

### Quantitative RT-PCR

Total RNA was extracted from melan-uw cells stably expressing either F374 or L374 variants of HA-SLC45A2 or HA-SLC35D3 as a control using the RNeasy Plus Kit (Qiagen), and 3 µg was reverse transcribed (RT) using the SuperScript III kit (Life Technologies). The resulting cDNA was used for qPCR. Primers were designed using Primer3 (http://bioinfo.ut.ee/primer3-0.4.0/) to span an exon-exon junction to avoid amplification of any contaminating genomic DNA, and were as follows: SLC45A2 (NM_016180.3) - F: CCCTGTACACTGTGCCCTTT and R: CTTCCCTCTCACGCTGTTGT. Reactions were prepared according to the manufacturer’s protocol using SYBR Select Master Mix (Invitrogen/ThermoFisher Scientific) and cycled on a Real-Time PCR System (Biorad). β-actin was used as an internal control and all reactions were run in triplicate. mRNA levels were quantified by calculating average 2^-ΔΔCt^ values, where Ct is the cycle number for the control and target transcript at the chosen threshold. ΔCt = Ct_target_ - Ct_β-actin_ was calculated by subtracting the average Ct of β-actin from the average Ct of the target transcript. The ΔCt_mean_ was calculated for the reference sample (L374) and ΔΔCt = ΔCt - ΔCt_mean_ was calculated by subtracting the ΔCt_mean_ from the ΔCt. The relative mRNA expression of samples compared to L374 samples was calculated by 2^-ΔΔCt^. Statistical significance was determined using one-way ANOVA with Dunnett’s test for multiple comparisons.

### Statistical Analyses

Statistical data are presented as mean ± SEM. Statistics were calculated in either GraphPad Prism or Microsoft Excel using one-way ANOVA or Student’s T test with correction where specified. Significant differences between control or experimental samples are indicated (****, p < 0.0001; ***, p < 0.001; **, p < 0.01; *, p < 0.05). Only p < 0.05 was considered as statistically significant.

## ACKNOWLEDGMENTS

The authors are grateful to Drs. Lynn Lamereux for generating *Slc45a2*^uw/uw^*Ink4a-Arf^-/-^* mice and Lynn Plowright for technical assistance in generating melan-uw cells from these mice, and to Dr. Sergio Grinstein for the gift of the LAMP-mCherry expression construct. We acknowledge funding from NIH grants R01 AR048155 (to MSM), R01 AR071382 (to EO and MSM), R01-AR066318 (to EO), and F32 AR062476 (to MKD) from the National Institute of Arthritis, Skin and Musculoskeletal Diseases and T32 GM007229 Training Program in Cell and Molecular Biology from the National Institute of General Medical Sciences (supporting LL and AJL), a National Science Foundation Graduate Research Fellowship (to KDH), and Wellcome Trust grant 108429/Z/15/Z (to EVS and DB).

## AUTHOR CONTRIBUTIONS

Linh Le designed and performed most of the experiments shown in the paper, designed and performed all of the statistical analyses, assembled all of the figures, and participated in drafting and editing the manuscript.

Iliana Escobar performed some of the experiments shown in the paper and repeated others, assembled drafts of the figures for these experiments, and edited the manuscript.

Tina Ho and Ariel Lefkovith designed and performed some of the experiments shown in the paper as well as experiments that formed the basis for other figures in the paper and that contributed to the quantification and statistical analyses of data shown. They both edited the manuscript.

Emily Latteri, Megan Dennis and Kirk Haltaufderhyde all designed and performed experiments that formed the basis of the project and that were included in the quantification and statistical analyses of data shown. All also edited the manuscript.

Elena Sviderskaya and Dorothy Bennett developed the immortalized melan-uw melanocyte cell lines and edited the manuscript.

Elena Oancea and Michael Marks conceived the project, contributed to the design of most of the experiments, assembled figures, and participated in drafting and editing the manuscript.

---

**Expanded View Figure EV1.**
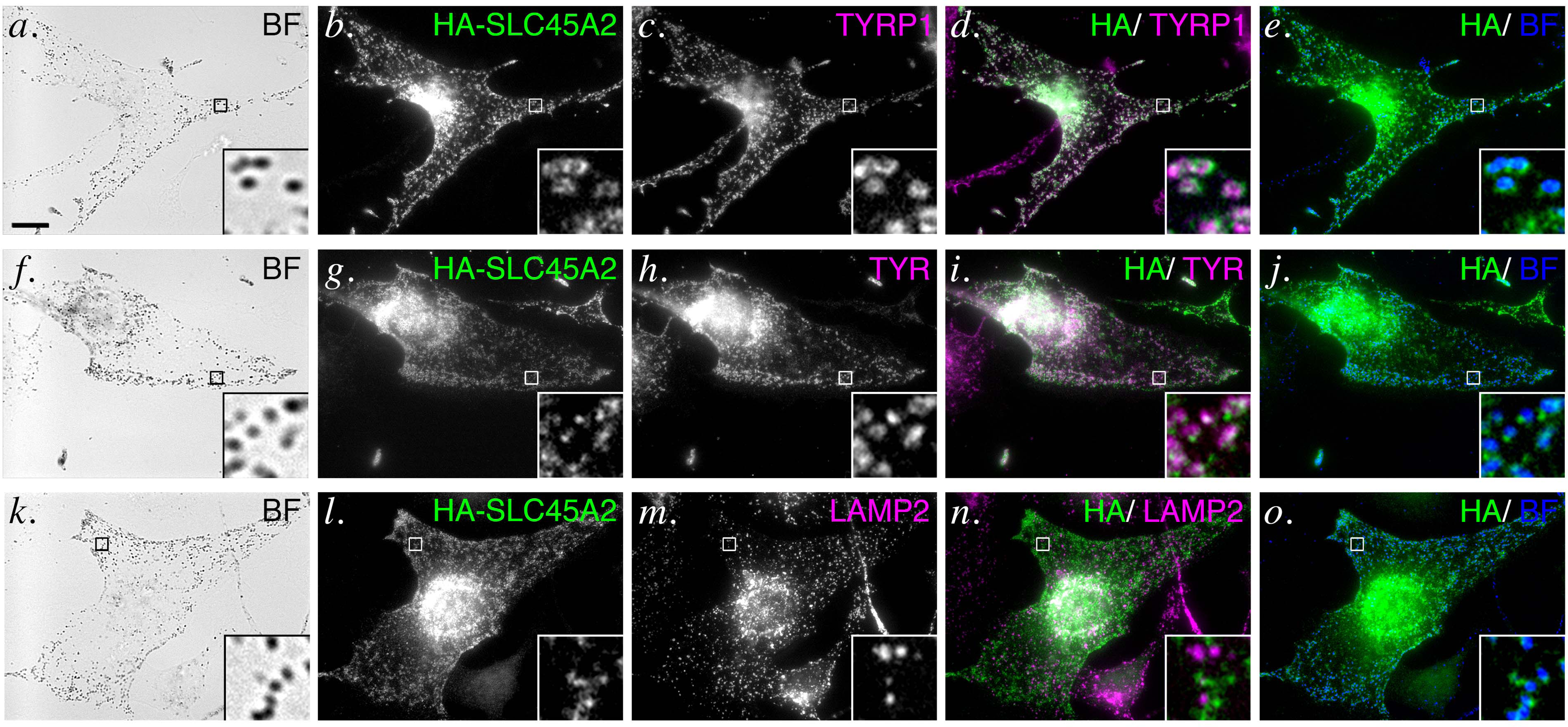
HA-SLC45A2 localizes to melanosomes upon transient transfection in WT melanocytes. Transiently transfected WT melan-Ink4a cells expressing HA-SLC45A2 were fixed, labeled for HA (green) and either TYRP1 (*a-e*, magenta), TYR (*f*-*j*, magenta) or LAMP2 (*k*-*o*, magenta), and analyzed by dIFM and by bright field microscopy. Shown are individual bright field (BF) or labeled panels (*a*-*c*, *f*-*h*, and *k*-*m*), merged dIFM images (*d*, *I*, *n*) or merged HA and bright field images (*e*, *j*, *o*; bright field is pseudocolored blue); boxed regions are magnified 7.5X in insets. Scale, 10 μm.

